# Functional connectivity between the entorhinal and posterior cingulate cortices associated with navigation impairment following path integration in at-genetic-risk Alzheimer’s disease

**DOI:** 10.1101/771170

**Authors:** Gillian Coughlan, Peter Zhukovsky, Vaisakh Puthusseryppady, Rachel Gillings, Anne-Marie Minihane, Donnie Cameron, Michael Hornberger

## Abstract

Navigation processes mediated selectively by the entorhinal cortex (EC) may be impaired in individuals with suspected preclinical Alzheimer’s disease (AD), but the clinical utility of navigation tests to detect such impairments remains to be established. In a sample of 64 individuals (32 e3e3 and 32 e3e4), we tested whether an existing paradigm, the Virtual Supermarket Test (VST), can reliably detect the presence or absence of the APOE e4 allele which accelerates amyloid plaque deposition in the brain. The present study assessed four major navigational processes that are subserved by functionally specialised cell groups located in AD vulnerable regions including the EC and examined the relationship between navigation process and regional functional connectivity (FC) given FC is a marker early AD-related tau seeding. While heading direction and spatial memory were unaffected by at-risk AD, clear altered navigational strategies following path integration were found on the VST in the e3e4 group. The APOE-sensitive VST measure correctly classified 77% of the APOE cohort. Including resting-state FC between the EC and the posterior cingulate cortex, a correlate of the path integration deficit in the APOE e4 group, the classification model increased the accuracy to 85%. Our findings show that at-genetic-risk AD selectively impairs path integration and biases self-reported spatial locations away from the centre and towards the boundary of a virtual environment. Importantly, this impairment is associated with reduced FC between the EC and the posterior cingulate cortex, which in turn informs the neurobiological mechanisms of at-genetic-risk of AD.

## INTRODUCTION

Late-onset Alzheimer’s disease (AD) is one of the biggest burdens to modern society with up to 50 million people living with the disease worldwide and no curative therapies to treat the underlying cause of the disease pathology (Nichols et al., 2019). Gold standard episodic memory tests fail to capture the first symptomatic manifestation of AD (Coughlan et al., 2018b; Jessen et al., 2014; Zimermann et al., 2019) and therefore, alternative diagnostic tools are urgently required.

Spatial navigation has emerged as a critical diagnostic tool in the clinical AD and mild cognitively impaired cohorts due to its sensitivity and specificity for AD pathophysiology (Hort et al., 2014, 2007; Howett et al., 2019; Laczó et al., 2009; Lester et al., 2017; Mokrisova et al., 2016; Pai and Jacobs, 2004; Serino and Riva, 2013; Tu et al., 2015; Vlček, K., & Laczó, 2014). For example, spatial navigation tasks, such as the Virtual Supermarket Test (VST) can distinguish AD from other dementias and have been implemented in many high-profile clinical trials as treatment outcome measures. At this stage however, it is uncertain if such tasks have any utility to detect impeding cognitive decline, particularly carriers of the Apolipoprotein E4 (APOE ε4) gene; a gene hypothesised to accelerate the β-amyloid plaque (Aβ) deposition and the neurodegenerative cascade in the brain years before the clinical onset of dementia (Jack et al., 2010; Mishra et al., 2018).

Extensive evidence on spatial navigation test to date shows that AD patients and individuals at an earlier stage of the disease spectrum, characterised by ‘mild cognitive impairment’ sustain widespread navigation deficits. AD patients show severe difficulty storing and retrieving a cognitive map of virtual environments (DeIpolyi et al., 2007; Jheng and Pai, 2009; Serino et al., 2015), consistent with significant cell loss in the hippocampus which is home to the place cells, the cellular building blocks of the human and rodent ‘cognitive map’ (Bird and Burgess, 2008; O’Keefe & Nadel, 1978). Navigation tests also show egocentric self-reference strategies are also lost in AD patients, due to various metabolic and volumetric changes that occur in the retrosplenial cortex, posterior cingulate cortex (PCC) and parietal cortex where head direction and border cells reside (Mokrisova et al., 2016; Pai and Yang, 2013; Pengas et al., 2010; Serino et al., 2015).

In the asymptomatic preclinical stage of AD, AD pathophysiology is relatively localised to the EC, while metabolic and resting-state functional changes occur in distant but connected regions, as the brain responses to early pathological growth deep within medial temporal lobe (Badhwar et al., 2017; Braak and Del Tredici, 2015; Chase, 2014). Nevertheless, preclinical experimental investigations to date primarily utilise tasks that tap into the integrity and function of EC, with the aim of identifying candidates most likely to covert to MCI or clinical AD due to the presence of early neuropathology (Fu et al., 2017; Howett et al., 2019; Kunz et al., 2015). Such studies reveal that errors in the entorhinal-mediated grid cells function correlates with altered navigation patterns during path integration (or self-motion) among at-genetic-risk APOE ε4 carriers who were otherwise asymptomatic for AD. The same pattern of altered navigation behavioural was replicated on the Sea Hero Quest game, and discriminated ε4 carriers from non-carriers, although no neural data was available (Coughlan et al., 2019). To add further support to the hypothesis that EC mediated navigation changes may represent an early cognitive marker for preclinical AD, transgenic rodent models show spatial memory deficits measured on the Morris water maze occur just before mature tau tangles spread beyond the entorhinal cortex (Fu et al., 2017).

Clinically then, the development of preclinical diagnostic tests relies on two key criteria. The first being that these tests measure and dissociate between different spatial processes, as each process relies on functionally separate cell including place cells, grid cells and boundary cells (Barry et al., 2006; Moser et al., 2008) located in different AD vulnerable brain regions. This dissociation is crucial when investigating AD-related spatial disorientation as early pathophysiology and resulting resting state functional and metabolic changes occur in very select brain regions, including the entorhinal cortex (EC), the precuneus and posterior cingulate cortex (Bott et al., 2016; Chase, 2014; Lazarczyk et al., 2012; Pengas et al., 2010). The second criterion is that spatial navigation tests are clinically feasible, which in turn would allow them to fulfil their purpose of being used as early screening measures in the mass population.

In the current study, we exploit growing evidance that EC mediated navigation process may well represent a marker for incipent/preclincal AD and took a novel approach to measure APOE ε4 related navigation changes using the well-validated AD diagnostic tool: ‘Virtual Supermarket Test (VST)’. We also tested navigation processes subserved by the PCC and precuneus as these AD vulnerable regions are key nodes in the human spatial navigation network. To that end, we measured four dissociable navigational processes subserved by functionally specialised cell groups located in the EC, the hippocampus, the PCC and the precuneus. As an additional measure, we assessed if subjective cognitive impairment, sometimes considered a first symptomatic manifestation of disease, accompanies loss (or intactness) of navigational process (Jessen et al., 2014). Finally, resting-state magnetic resonance imaging (MRI) data was collected to measured functional connectivity (FC) between these regions with two aims: i) to provide insight into the functional neurobiology of at-genetic-risk of AD and ii) to identify a neural correlate for navigation impairment(s) sustained by the APOE ε4 group. On a behavioural level, we hypothesis that ε4 carriers would show one or more navigation impairments on the VST. On a neural level, we hypothesis that the navigation impairments will be correlate with functional connectivity between the EC and the PCC/precuneus.

## MATERIALS AND METHODS

### Participants

We recruited 150 participants between 50 and 75 years of age (M=61.92, SD=6.72) to participate in a research study at the University of East Anglia. Written consent was obtained from all participants and ethical approval was obtained from Faculty of Medicine and Health Sciences Ethics Committee at the University of East Anglia, Reference FMH/2016/2017–11. All participants were pre-screened over the phone for a history of psychiatric or neurological disease, history of substance dependence disorder or any significant relevant comorbidity, such as clinical depression or anxiety. History of antidepressant treatment with serotonin reuptake inhibitor drugs was retrospectivity obtained. Participants receiving anti-depression or anti-anxiety medication at the time of screening, were excluded as both conditions impact cognitive performance and self-report cognitive decline (Halahakoon, Lewis, 2019). Moreover, only participants with normal or corrected-to-normal vision were retained due to the nature of the VR task. As family history was assessed based on remembered number of parents (0, 1 or 2) with dementia, this was a retrospective account of which participants express uncertainty, particularly around the type of dementia and thus not included in the main analysis. Finally, saliva samples were collected via buccal swab from those who passed this screening and APOE genotype status was determined.

As just 25% of the population carry an APOE ε4 allele (23% APOE ε3ε4, 2% APOE ε4ε4; Corbo and Scacchi, 1999; Liu *et al.*, 2013), all ε4 allele carriers in our sample were retained for the experiment. A subset of the ε3ε3 carriers that form the majority of the population (60%) to match the ε3ε4 risk group for age and sex were included (see Table S1 for group background characteristics), which brought us to a final sample size of 64 including 32 ε3ε3 carriers and 32 ε3ε4 carriers) all of whom underwent cognitive testing. Twenty ε3ε3 carriers and 20 ε3ε4 carriers also underwent structural and functional MRI. We did not include a third genetic subgroup of homozygous APOE-ε4 carriers from the tested cohort, because they were too rare (*n* = 4). APOE ε2carriers (15% of the UK population) were also excluded as it is unclear how the ε2 allele acts on cognitive performance or the further development of AD. One ε3ε3 participant did not complete the scan due to distress and their data were excluded from the analysis. Two additional participants (one ε3ε3, one ε3ε4 carrier) who completed the MRI stage of the study were removed due to a software error that led to severe artefacts in the resting-state fMRI data. After these exclusions, MRI data on 37 out of 64 (58%) of the cognitively tested participants remained.

### Paradigm overview

The VST is a sensitive and specific measure for differentiating AD from other dementia types (Coughlan et al., 2018a; Tu et al., 2017, 2015) and taps into i) translation from allocentric to egocentric self-reference navigation also term egocentric orientation; ii) short-term spatial memory; iii) heading direction and iv) central (vs boundary) based navigation preference. In brief, an iPad 9.7 (Apple Inc,) is used to show participants 7-14-second video clips of a moving shopping trolley in a virtual reality supermarket from the first-person perspective (Figure 1 A-C). The absence of landmarks in the supermarket aims to ensure the test taps into EC-grid cell dependent strategies rather than striatal-mediated landmark-based navigation. Once the video clip stops, participants indicate in real-life the direction of their starting point (egocentric orientation; Figure 1 D). In a second step, participants indicate their finishing location (short-term spatial memory; Figure 1 E) and heading direction on a VST map. We extended our VST paradigm to a fourth spatial measure based on evidence of an entorhinal-mediated bias or tendency to navigate towards environmental boundaries during path integration in at-genetic-risk AD. Specifically, we recorded the number of responses in the centre and boundary space of the supermarket map to produce a central navigation preference measure (Figure 1 F; see supplementary text for more information on the central navigation preference measure).

**Figure 1.**
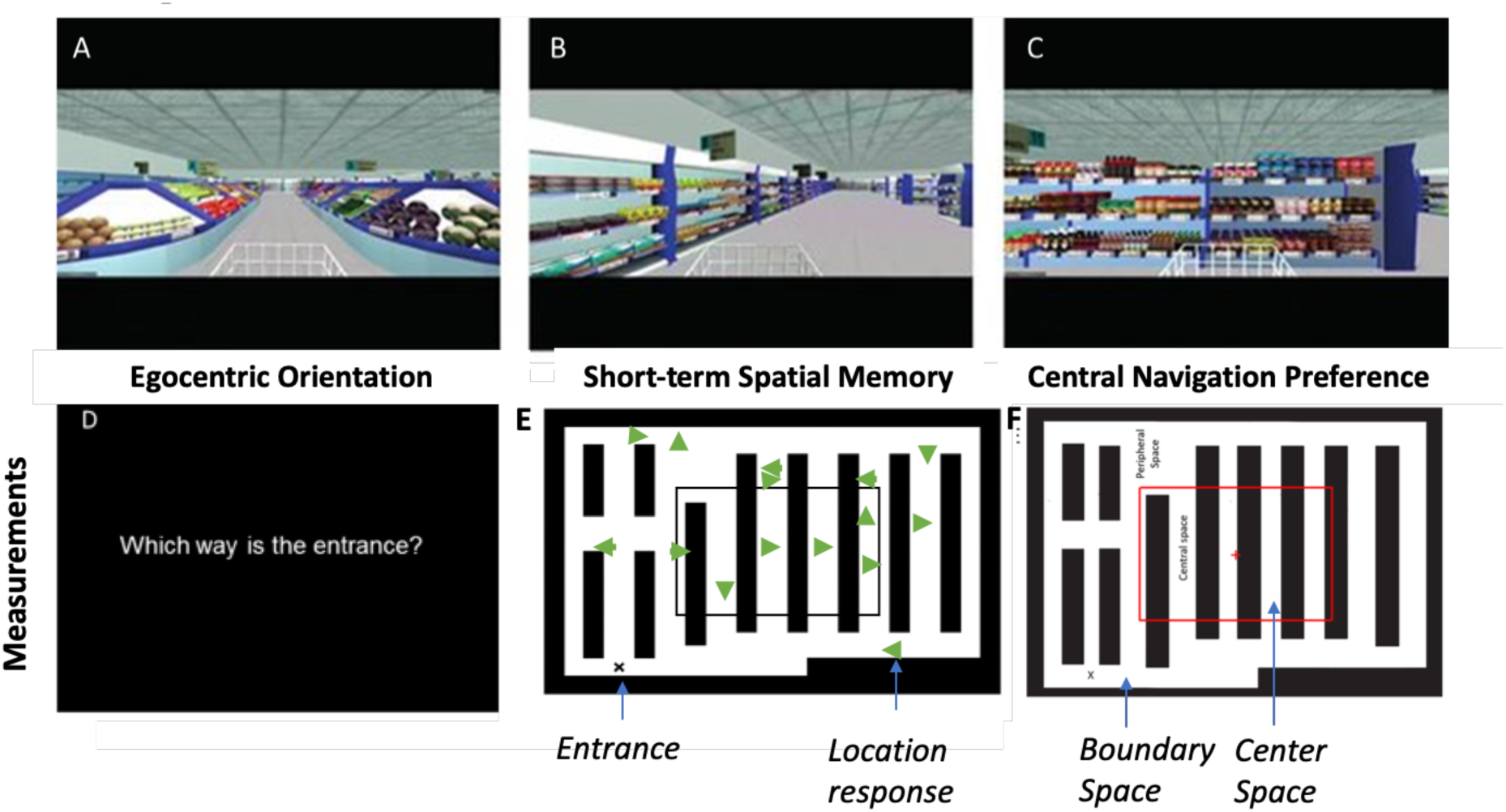
Spatial orientation was assessed using an ecological virtual supermarket environment. The layout of the virtual environment did not include any notable landmarks. An iPad 9.7 (Apple Inc., etc) was used to show participants 7-14-second video clips of a moving shopping trolley. All trials began at the same locationin the supermarket, but followed different routes to reach a different end point in each trial (**A**). Videos were presented from a first person perspective and participants were taken to a set locations while making a series of 90 degree turns (**B**). Once the video clip stopped (**C**), participants indicate the real-life direction of their starting point (**D**). Immediately following, participants indicate their finishing location (short-term spatial memory) and heading direction on a VST map (**E**). Number of location responses made in the center space and boundary spaces were recorded (**F**)

### Neuropsychological assessment

The aim of the current study was to assess the impact of APOE genotype independent of, and prior to, AD symptomology. The Addenbrooke’s cognitive examination (ACE-III) was used to detect cognitive impairment associated with AD (Matias-Guiu et al., 2017). Only participants who scored in the normal range (ACE-III>88) were retained. The Rey–Osterrieth Complex Figure Test (RCFT; with 3-min delayed recall) and the Four Mountains test were used as secondary screening measures to assess any differences between genetic groups (Chan et al., 2016; Shin et al., 2006).

### Subjective cognitive change assessment

Subjective cognitive decline was evaluated to identify decline in self-perceived episodic memory and executive function compared to 5 years before testing. In prior work, subjective memory concerns have been identified in asymptomatic familial AD carriers, and concerns are seemly predictive of faster rates of memory decline (Samieri et al., 2014; Weston et al., 2018). The presence of subjective cognitive concerns is also related to abnormal changes in Aβ and tau biomarkers in APOE ε4 carriers (Risacher et al., 2015) and is thus considered important for early detection. Here, we measure this using the Cognitive Change Index (CCI; Rattanabannakit et al., 2016) that consists of 20 questions relating to the perceived decline. Responses are given on a five-point scale ranging from 1 = “normal ability” to 5 = “severe problem”, with higher scores indicating larger concerns.

### APOE Genotyping

Please refer to Coughlan et al 2019, for a description on the APOE genotyping method used for the present study.

### Functional MRI acquisition

Structural and functional MRI data for 40 participants (20 ε3ε3 carriers and 20 ε3ε4 carriers) was obtained using a 3 tesla Discovery 750w widebore system (GE Healthcare, Milwaukee, WI, USA) with a 12-channel phased-array head coil for signal reception. After localisers, T_1_-weighted (T_1_w) structural data was acquired using a whole-head 3D inversion-recovery fast spoiled gradient recalled echo (IR-FSPGR) sequence with the following parameters: repetition time = 7.7 ms; echo time = 3.1 ms; inversion time = 400 ms; field-of-view = 256 mm; acquired matrix = 256 × 256; 200 sagittal sections of 1 mm thickness; flip angle = 11°; and an ASSET acceleration factor of 2 in the phase-encoding direction. Furthermore, a 3D T_2_-weighted fluid attenuated inversion recovery (T_2_w FLAIR) sequence was prescribed as follows: repetition time = 4,800 ms; echo time = 129 ms; inversion time = 1,462 ms; field-of-view = 256 mm; acquired matrix = 256 × 256; 182 sagittal sections of 1 mm thickness; flip angle = 90°; an ARC acceleration factor of 2 in the phase-encoding direction; and a ‘HyperSense’ compressed sensing subsampling factor of 2. Functional images were acquired using a gradient echo echo-planar imaging sequence with the following parameters: repetition time = 3,500 ms; echo time = 30 ms; field-of-view = 240 mm; acquired matrix = 96 × 96, reconstructed to 128 × 128; 42 axial slices of 3.5 mm thickness; flip angle = 80°; and an ASSET acceleration factor of 2 in the phase-encoding direction. The fMRI time series consisted of 200 images, and the total acquisition time was 11 minutes 54 seconds. During functional runs, subjects were required to not fall asleep and keep alert with their eyes closed for 10 min. To avoid the effect of participants employing specific strategies to maintain alertness (e.g. reminiscing or counting scan number), participants were instructed not to think about anything in particular. Prior to analyses, all participant scans were visually inspected for significant head movements and artefacts. Please see supplementary materials for pre-processing structural and functional MR images.

### Statistical analysis

Statistical analysis was performed using SPSS (v25.0), FSL (v6.0.0), MATLAB (MathWorks, R2018a), Octave (v4.4.1) and FreeSurfer (v11.4.2). Chi square and independent two tailed *t*-tests were used to test the significance of any demographic or neuropsychological assessment differences between the genetic groups. All group comparisons were conducted using the same general linear model including APOE, as main predictor of interest. Age and sex as covariates given their strong effect on brain function and volume, navigation performance and AD development (Coutrot et al., 2018; Ferretti et al., 2018; Lester et al., 2017; Neu et al., 2017). All scripts ran in FSL, MATLAB, Octave and FreeSurfer are available upon request.

Performance on each sub-measure of the VST and the CCI were entered a univariate analysis of covariance. Associations between VST sub-measured and CCI was tested using partial Pearson correlation in SPSS. Voxel-based morphometry (VBM) was conducted on whole-brain T_1_weighed scans, using the VBM toolbox in FSL to confirm no grey and matter structural differences between the genetic groups (Douaud et al., 2007; Good et al., 2001). FreeSurfer was used to segment and parcellate whole-brain T_1_weighed images to provide a volumetric measure for the apriori selected anatomical ROIs. This was done in favour of VBM masking. Total intracranial volume was included as a covariate. FC between pairs of the four ROIs were analysed by extracting the first eigenvector from the BOLD timeseries for each ROI, and each single participant, using ‘fslmeants’. A total of 195 of the 200 functional timepoints for each ROI were retained for analysis. All functional network modelling with timecourse data was carried out using FSLNets v0.6 so that the functional connectivity results were family wise error (FWE) corrected for multiple comparisons. After computing the subject-specific 6^nodes^ × 6^nodes^ connectivity matrix, direct and ridge regularised partial correlations were calculated between all pairs of ROIs, where direct correlations are correlations between two ROIs, controlling for the effect of all other ROI-ROI correlations. The resulting Pearson correlation coefficients were converted to z scores via Fisher’s transformation to test the significance of any functional connectivity differences between the genetic groups (Smith et al., 2011). All functional analyses were carried out in MNI standard space. Significance testing for all MRI differences was conducted using voxel-wise general linear modelling by employing the threshold-free cluster enhancement (TFCE) method (Smith and Nichols, 2009). The TFCE produces voxel-wise P-values via 5,000 permutation-based non-parametric testing (Nichols and Holmes, 2001).

## RESULTS

### Neuropsychological assessment

As expected, no differences between the two genetic groups were present on the neuropsychological assessment, which confirmed that the impact of APOE genotype prior to clinically detectable MCI/AD symptomology could be measured (also see SI Table 1 for secondary demographics).

**Table 1.** Primary demographic and neuropsychological profile.

| | $\epsilon 3\epsilon 3$ carrier (n=32) | $\epsilon 3\epsilon 4$ carrier (n=32) | *P* value |
| --- | --- | --- | --- |
| Age (years) |  |  |  |
| Mean (SD) | 62.24 (5.32) | 62.19 (5.58) | - |
| Sex |  |  |  |
| Male | 17 | 22 | - |
| Female | 15 | 10 | - |
| ACE | 94.47 (3.83) | 92.88 (3.78) | .12 (F=2.49) |
| FMT | 10.22 (2.91) | 9.47 (1.23) | .49 (F=.484) |
| ROCT |  |  |  |
| Recall | 22.92 (2.77) | 17.66 (4.95) | .06 (F=2.061) |
| Copy | 33.77 (6.36) | 32.12 (2.67) | .57 (F=1.287) |

Primary demographic and neuropsychological characteristics of the genetic groups (Independent sample t-test, two-tailed). ACE= Addenbrooke's cognitive examination. FMT = Four mountains test. ROCT = The Rey–Osterrieth complex figure. Recall administered three minutes after copy.

### Spatial navigation assessment

Heading direction (F = 1.87., *P =* 0.07) and short-term spatial memory (F=.572, *P =* .455) were unaffected by genotype and thus, we concluded group differences on other VST sub-measures could not be accounted for by differences in short-term spatial memory ability. Egocentric orientation was significantly different between genetic groups, as ε3ε4 participants made fewer correct responses compared with ε3ε3 participants (F = 4.18; *P =* 0.04). Central navigation preference was also significantly different between the groups (F = 12.45, *P* < 0.005), with ε3ε3 participants favouring more central responses and ε3ε4 carriers favouring more boundary responses (see Figure 2 A-C; Table 2 for mean values).

**Figure 2.**
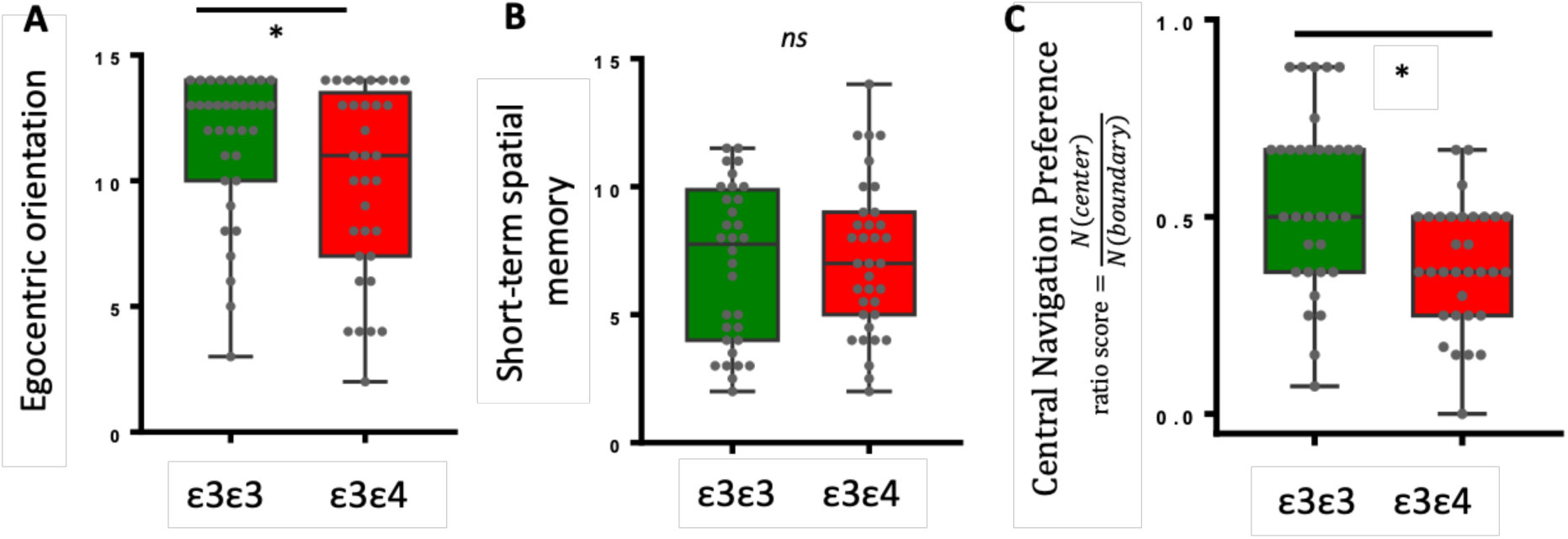
ANCOVA showed an effect of genotype on egocentric orientation and central navigation preference. No effect of genotype was present on short-term spatial memory.

**Table 2.** Effect of genotype on the VST spatial orientation paradigm.

|  | *Mean (SD)* | *F* | *P* value |
| --- | --- | --- | --- |
| Egocentric orientation |  |  |  |
| $\epsilon 3\epsilon 3$ | 12.01 (2.3) | 4.21 | .042 |
| $\epsilon 3\epsilon 4$ | 10.94 (3.7) | | |
| Heading direction |  |  |  |
| $\epsilon 3\epsilon 3$ | 11.92 (2.7) | .799 | .375 |
| $\epsilon 3\epsilon 4$ | 11.31 (3.0) | | |
| Allocentric memory |  |  |  |
| $\epsilon 3\epsilon 3$ | 7.43 (2.7) | .014 | .907 |
| $\epsilon 3\epsilon 4$ | 7.34 (3.0) | | |
| Central vs boundary preference |  |  |  |
| $\epsilon 3\epsilon 3$ | .57 (.21) | 12.45 | < 0.005 |
| $\epsilon 3\epsilon 4$ | .38 (.14) | | |

ANCOVA with age and sex as covariates testing the difference of egocentric orientation, heading orientation, allocentric memory and central preference.

**Table 3.** Summary of ε4 related cognitive disturbances.

| | $\epsilon 4$ related impairment | Neural correlate identified |
| --- | --- | --- |
| *Measure* |  |  |
| Egocentric orientation | Yes | Not identified |
| Heading direction | No | Left-EC- PCC |
| Short-term spatial memory | No | Not identified |
| Central navigation preference | Yes | Right EC- PCC |
| Subjective episodic memory decline | Yes | Right EC- PCC |
| Subjective executive function decline | Yes | Not identified |

Adjusted for age and sex.

### Subjective cognitive change assessment

Next, we examined the significance of any differences on self-reported cognitive decline (within the last 5 years) between the genetic groups. ε3ε3 participants reported less decline on both episodic memory (F=5.24 *p*=.026) and executive function (F=5.92 *P*=.018) measured by the self-report CCI. Thus, we then sought to test the associations between navigation performance on the VST and CCI scores. Heading orientation, short-term spatial memory and central navigation preference were not significantly associated with CCI scores. Only egocentric orientation was associated with self-reported decline in executive function (r=-.347, *p*=.008), revealing that better egocentric performance is related to less subjective decline.

### Volumetric and/or functional connectivity

Having clarified the behavioural manifestation of ε4-related navigation impairment on the VST, we sought to investigate 1) the significance of any volumetric differences and/or functional connectivity changes between genetic groups and 2) if a neural correlate(s) for e4-related navigation impairment on the VST could be identified. A VBM analysis, with a whole brain mask revealed no significant grey matter volumetric differences between the groups (*p =* 0.18). The mean volumes of pre-selected ROIs (right/left hippocampus, right/left EC, PCC, Precuneus; see Figure 3A) also did not differ between groups (see supplementary table 1 for mean ROI volumetric values in both genetic groups). Next, we examined FC between selected ROIs, and investigated differences in connectivity strength between the genetic groups. Right EC and PCC connectivity was significantly lower in ε3ε4s relative to ε3ε3s (t=-2.608; uncorrected p=.01; corrected p=.03; 95%CI [-.426 -.053]; r_s_ = .171, F=6.80, p=098). Partial correlations (ridge regularised) were significantly lower in ε3ε4 carriers, even after multiple comparison correction (Fstat, pstat for the overall model, T stat for the APOE effect, pstat uncorrected, *P*_*FWE*_ *=* 0.027) whereas full correlations were only significantly lower at the uncorrected level (uncorrected *P =*0.017; *P*_*FWE*_ *=* 0.157). Trend differences in the opposite direction were observed between the precuneus and the PCC, with higher FC between these regions in ε3ε4s compared to ε3ε3s (t=-2.225; uncorrected p=.03; corrected p=.06; 95%CI=[.009 .214];; r_s_ = .228; P=035). Note Figure 3 B represents differences between ε3ε3 vs ε3ε4 on the connectivity correlation matrix that illustrates the significance of differences between ROI-ROI Pearson correlation values between groups. *red circle* = ε4 related reduced connectivity, *green circle* = ε4-related increased connectivity). See supplementary materials for the ε3ε3 and ε3ε4 connectivity matrices.

**Figure 3.**
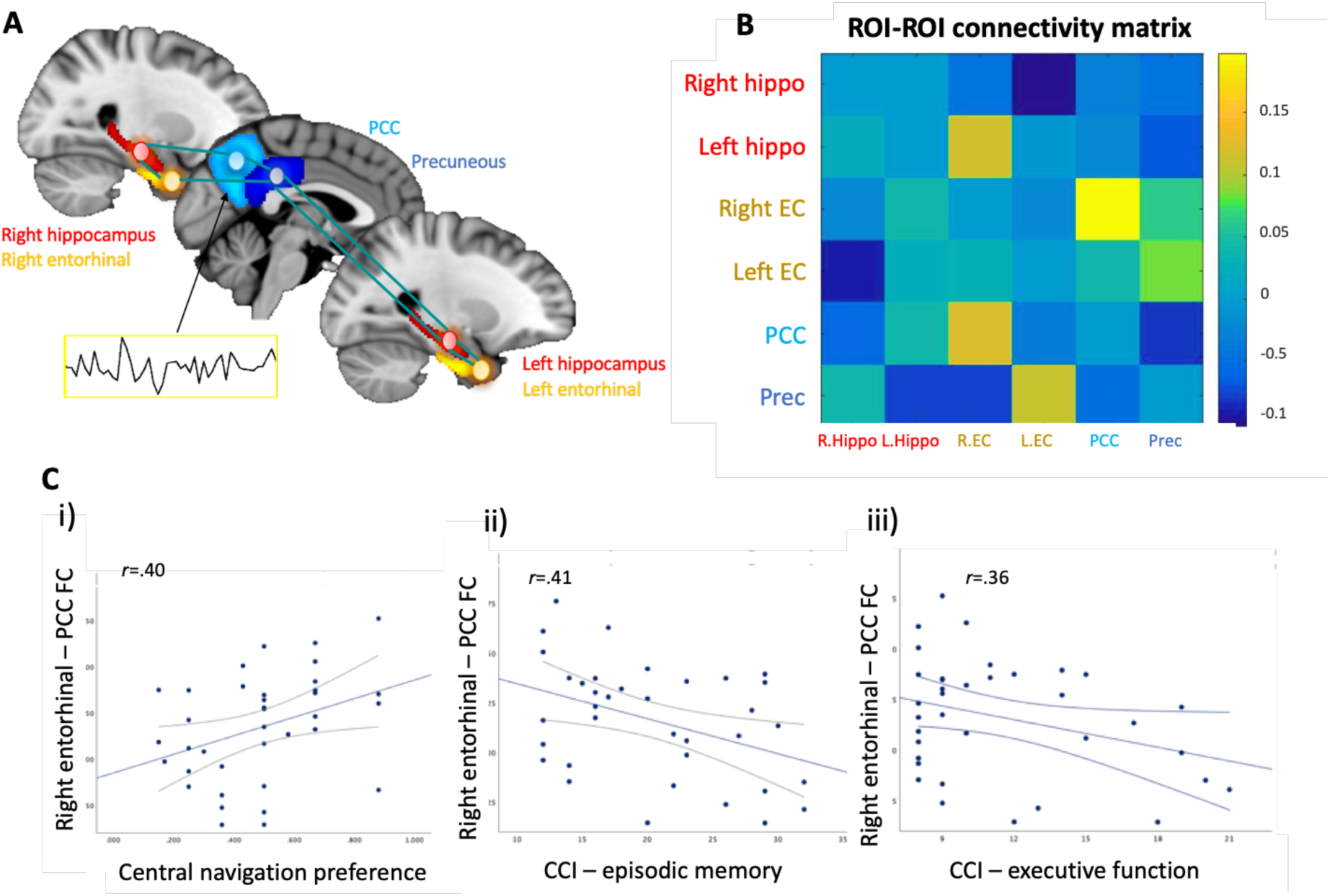
A APOE-dependent correlations in functional connectivity between selected ROIs: *right/left hippocampus, right/left EC, PCC, Precuneus*. **B** The 6node × 6node network matrix of correlation coefficients represents connectivity strength between nodal pairs in a dual regression to test two-group subject difference on subject specific nodal pair connectivity. **C** Linear regression models adjusted for age and sex show that i) right EC-PCC connectivity predicts central navigation preference ii) right EC-PCC connectivity predicts degree of perceived episodic memory decline and iii) right EC-PCC connectivity does not significantly predict the degree of perceived executive function decline.

**Figure 4.**
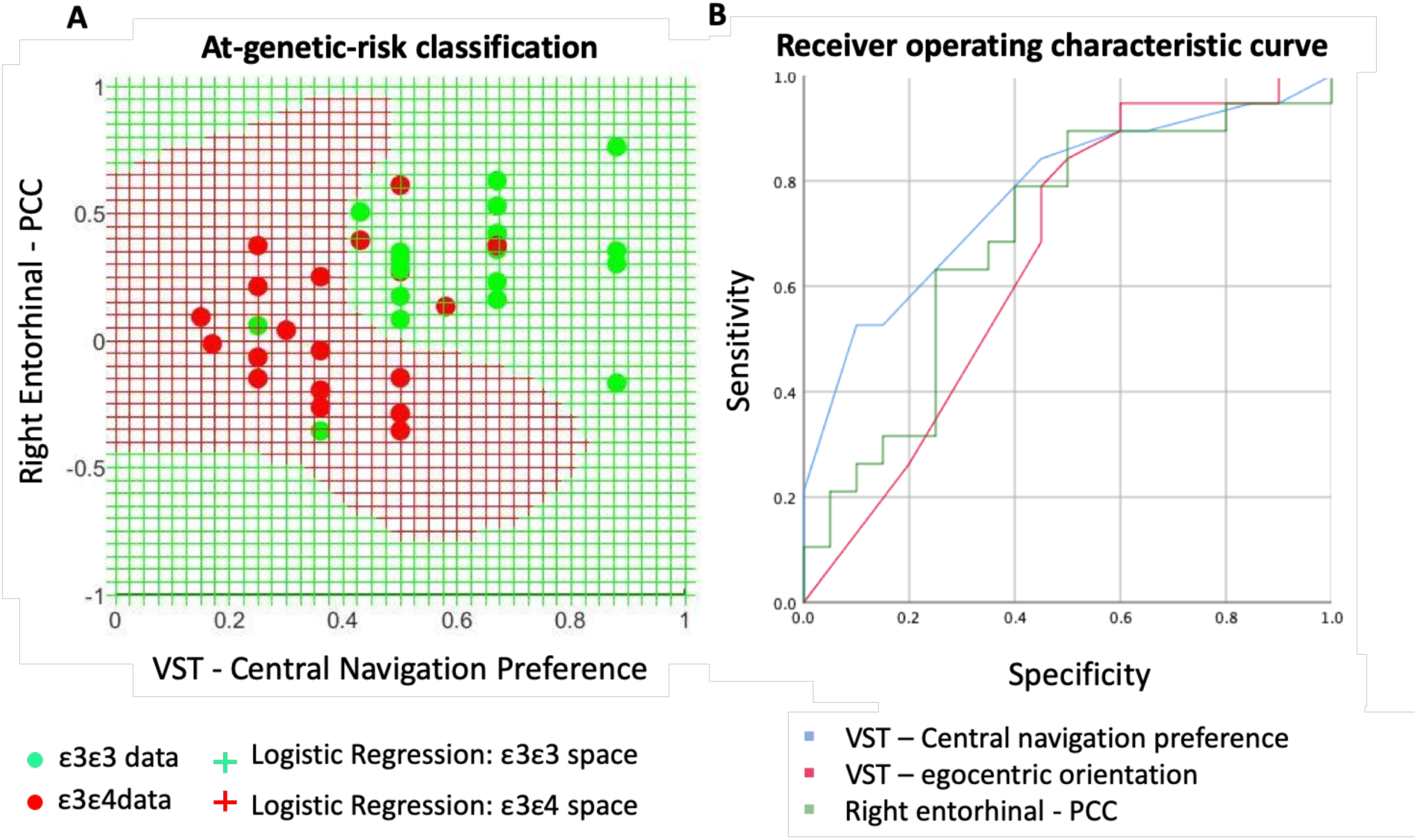
ROC curves for right EC– PCC functional connectivity strength (green line) and VST cognitive measures central preference (blue) and egocentric orientation (red) predicting variants of the APOE genotype. **A** Logistic regression indicated that the regression model based on function connectivity and VST cognitive predictors was statistically significant. **B** Area under the curve (AUC) values indicated EC-PCC and egocentric orientation had a similar level of diagnostic accuracy, while central preference had the best accuracy of the three predictors.

### Functional connectivity and ε4 sensitive navigation processes

Having determined altered functional connectivity changes in the EC, PCC and precuneus between genetic groups, connectivity strength between each ROI pair was correlated the ε4 sensitive VST measures: egocentric orientation and central navigation preference. This analysis was again performed using FSLNets to correct for multiple comparisons, meaning only robust correlations could be reported as significant. Functional correlates were identified for central navigation preference but not egocentric orientation. Specifically, right EC-PCC connectivity strength negatively correlated with central navigation preference when full correlations (r=0.40, *P*_*FWE*_ *=*0.018) were used as a connectivity metric (Figure 3. Ci). See supplementary materials for investigations on the neural correlates of the two additional VST sub-measures (short term-spatial memory and heading direction).

### Functional connectivity and subjective cognitive decline

Based on the e4-related changes on self-reported cognitive decline, we then measured functional connectivity with CCI scores. Connectivity strength between the right EC – PCC was negatively correlated with subjective decline in episodic memory (r=-.407 *P*=.017), but not executive function (r=-.336 P=.052; Figure 3 C).

### Classifying genetic groups based on VST and functional connectivity

Although no neuro-functional correlate was identified for egocentric orientation or subjective executive function decline, the neuro-functional correlate of central navigation preference and subjective episodic memory decline overlapped. Thus, as a final step, we tested its clinical utility to classify at-genetic-risk AD. In the first instance, we did not include functional connectivity and subjective decline measures, as our primary aim was to test the diagnostic value of the VST for at-genetic-risk AD. Thus, the first logistic regression model entered aimed to classify ε3ε3 and ε3ε4 carriers based on central navigation preference and egocentric orientation measure. This statistically significant (x^2^(2) 20.22, *P* < .001) and correctly classified 77.4% of the overall cohort (n=64). The percentage of classification was equal across ε4 carriers and non-carriers (Figure 5A). We then included the right EC –PCC measure to weigh the utility of including a neuro-functional marker to improve the classification. Note the sample size dropped to 37 with the inclusion of MRI measures. As expected, the regression model was statistically significant, x^2^(3) 16.85, *P* < .001) and classification accuracy shifted from 77.4% to 85%. Specifically, the model correctly classified 82.3% of ε3ε3 carriers and 88.3% of the ε3ε4 carriers (Figure 5 A). The log odds units presented are the values for the logistic regression equation for predicting APOE status from the three independent variables. The prediction equation is:

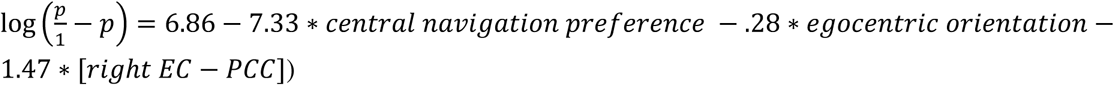

ROC curves were computed with these three predictors. Area under the curve (AUC) values indicated right EC-PCC connectivity (AUC .702, SE .092) and the egocentric task (AUC .659, SE .098) had a similar level of diagnostic accuracy. Central navigation preference showed the best accuracy of the three predictors (AUC .810, SE .073; Figure 5 B).

## DISCUSSION

APOE ε4 is the strongest genetic risk factor for late-onset AD. Yet, whether ε4-related cognitive changes are clinically detectable is unknown. Our results show that i) select ε4-related navigational changes following path integration occur alongside subjective concerns of cognitive decline ii) ε4-related navigational changes and self-report episodic memory concerns share the same neurobiological correlate iii) and the classification accuracy of the ε4-senstive VST sub-measures, when coupled with the right EC - PCC resting-state FC, reaches 85%, which in turn provides a springboard for the development of a simple multimodal framework for at-genetic-risk AD.

Significant differences between the genetic groups were found in two out of four of the VST spatial sub-measures: Egocentric orientation and central navigation preference. Egocentric orientation requires participants to form an accurate representation of the supermarket environment during self-motion, and then integrate this representation at the finishing location to produce an accurate directional representation of the starting point. ε4 carriers demonstrated significantly more difficulty identifying their starting point, suggesting ε4-related problems integrating allocentric-egocentric frames. Although short-term forgetting could explain this effect, the ε4 group showed no impairments on the spatial memory control measures, compared to the non-carrier group (i.e. the VST short-term spatial memory task and the four mountains task), making a memory-based causation unlikely. The central navigation preference adopted for this study measures the number of allocentric location responses in the central vs the boundary area of the virtual supermarket following the path integration task. This measure then provides a means of dissociating between central vs the boundary responses preferences (or biases). The most striking ε4-related behavioural discrepancy appeared on this measure, as ε4 carriers showed a strong responses bias towards the boundary, compared to non-carriers.

Behaviourally, the ε4-related bias for reduced central navigation preferences is consistent with entorhinal-mediated navigation pattern changes *during* path integration observed on two other experimental navigation tasks (Kunz et al., 2015; please see Coughlan et al., for an account on grid-cell representations underlying boundary-based navigational discrepancies in genetic at-risk individuals of AD). This is the first time ε4-related boundary biases were found *following* path integration, however, and although no neural FC correlate emerged for ε4-related egocentric orientation deficit, reduced FC between the right EC - PCC emerged as a significant neural substrate for reduced central navigation preferences in the at-genetic-risk group. Right EC – PCC FC also predicted the degree of episodic memory decline, as reported on the CCI (see Contreras et al., 2017 for more information on the CCI).

APOE ε4 carriers with self-assessed cognitive concerns have previously been shown to the have worse memory but this is the first study to show APOE ε4 carriers with self-assessed memory and executive function decline also show navigation discrepancies, and that navigation discrepancies and perceived episodic memory decline are mediated be the functional connectivity strength of neural pathways between the EC and PCC. Subjective memory decline with unimpaired performance on standard medical tests has been considered a first symptomatic manifestation of AD and is believed to be predictive of early AB accumulation (Jessen et al., 2014; Ngandu, T. et al. 2007; Dik, M. G. et a 2001). Considering subjective cognitive decline and subtle navigation deficits appear on our tasks (that at the same time does not reach the level of objective impairment required for the prodromal AD diagnosis), we conclude that subjective complaints may well contribute to a sensitive and specific diagnosis of preclinical AD, although the relevance of subjective concerns for clinical practice is outside the boundaries of this study. Moreover, it is difficult to say if navigation changes precede subjective cognitive deficits or vice versa.

The role of reduced EC – PCC functional connectivity in preclinical AD discovered here is not be surprising, as typically intracellular tau projects from the hippocampus and surrounding areas, to the PCC in the first stages of disease (Belloy et al., 2019; Jacobs et al., 2018). This spread is consistent with animal models that show in amyloid positive rodents, tau pathology propagation begins in the EC before spreading to the parietal cortex (Ahmed et al., 2014; Khan et al., 2013). Therefore, this pattern of neuropathological projection may well help explain the reduced functional connectivity in the at-genetic-risk group, as well as the impeded translation of the allocentric or egocentric coordination system, given that the allocentric system relies on entorhinal-hippocampus regions and the egocentric system relies on parietal/PCC regions. We cannot say for sure however, as the egocentric orientation measure did not correlate with the FC strength between the EC and PCC. It may be the that egocentric orientation changes are underpinned by functional changes between regions not examined in the present study, for example, frontal lobe regions of the brain. This is certainly possible, given the widespread APOE4-related functional changes reported in the literature to date (Badhwar et al., 2017). Moreover, egocentric orientation is associated with perceived executive function ability making it more likely that the egocentric task shared some dependencies on the frontal lobes. We also observed increased PCC-precuneous connectivity in the genetic-risk group. This effect may be best explained by animal models that demonstrate moderate levels of Aβ in the brain can enhance FC due to compensatory mechanisms, which would explain why increased connectivity strength in intrinsic brain networks is commonly found in APOE ε4 cohorts (Badhwar et al., 2017; Chase, 2014; Machulda et al., 2011).

Despite our results largely supporting current theories of preclinical AD models, our results has several limitations. Firstly, our sample size fell from sixty-four to thirty-seven when investigation the neural correlates of APOE4 related navigation impairment on the VS. Thus, could help explain the fact that we could not identify a neural correlate for egocentric navigation and executive function, both of which were significant association with each other. Alternatively, it may be that functional connectivity between the four AD vulnerable regions simply does not exert a significant effect on these cognitive processes. This is possible as egocentric navigation has been strongly linked to the structural integrity of the RSC. We cannot rule out the possibility that the boundary-driven navigation behavioural following PI is caused be another neural mechanism and/or the fact that landmarks, although intentionally hidden in the VST, may influence to draw toward to border in the APOE4 group Future studies might also consider applying graph theory to a larger sample size; examining the neural characteristics of the AD-vulnerable spatial network and investigating overall network efficiency in terms of information passing. In addition, we would recommend a more robust assessment of AD risk for future studies, including AB and tau evaluation from cerebrospinal fluid and in the brain, which would in turn give greater validation to the VST as a diagnostic tool of underlying AD pathology. Alternatively, we would longitudinally follow-up of participants to confirm whether the observed changes are indeed predictive of future development of clinical AD is required.

In conclusion, we have shown a distinctive association between navigational deficits and altered FC in three key nodes of the spatial navigation network. As recent clinical trials of disease-modifying agents in Alzheimer’s disease have failed (Sevigny *et al.*, 2016), the addition of simple multimodal diagnostic models for at-risk AD could facilitate the shift from treatment to prevention, allowing targeted interventions with potential pharmaceutical and non-pharmaceutical neuroprotective compounds, in the preclinical stage of AD (Dubois et al., 2014; Reiman et al., 2015). If our results in cognitively normal middle-aged APOE e4 carriers do reflect a very early functional change in the course of AD progression, then the VST could also be a useful means of enrolling individuals in future clinical trials as early as possible.

## Data Availability

Link will be provided https://osf.io

## Acknowledgements

We would like to thank Dr Sicong Tu who co-developed the virtual supermarket test. We also wish to extend a thank you to all participants who devoted their time to this research.

---

**Supplementary Table 1.** The effect of APOE on regional ROI volume.

|  | Mean (SD) | P value |
| --- | --- | --- |
| Right entorhinal |  |  |
| $\epsilon 3\epsilon 3$ | 1872.8 (355.5) | *ns* |
| $\epsilon 3\epsilon 4$ | 1674.4 (274.9) | |
| Left entorhinal |  |  |
| $\epsilon 3\epsilon 3$ | 1935.0 (326.0) | *ns* |
| $\epsilon 3\epsilon 4$ | 1975.2 (398.5) | |
| Left PCC |  |  |
| $\epsilon 3\epsilon 3$ | 2889.8 (328.3) | *ns* |
| $\epsilon 3\epsilon 4$ | 3087.1 (508.7) | |
| Right PCC |  |  |
| $\epsilon 3\epsilon 3$ | 3000.1 (447.1) | *ns* |
| $\epsilon 3\epsilon 4$ | 3016.8 (407.2) | |
| Right precuneus |  |  |
| $\epsilon 3\epsilon 3$ | 9645.8 (909.9) | *ns* |
| $\epsilon 3\epsilon 4$ | 9875.0 (1307.6) | |
| Left precuneus |  |  |
| $\epsilon 3\epsilon 3$ | 9334.2 (818.8) | *ns* |
| $\epsilon 3\epsilon 4$ | 9317.8 (1067.2) | |
| Right hippo |  |  |
| ε3ε3 | 3502.9 (325.5) | *ns* |
| ε3ε4 | 3483.4 (422.5) |  |
| Left hippo |  |  |
| ε3ε3 | 3416.7 (358.0) | *ns* |
| ε3ε4 | 3367.8 (390.1) |  |
| ANCOVA with age and sex as covariates testing the difference regional brain volume |  |  |

